# Mapping cell structure across scales by fusing protein images and interactions

**DOI:** 10.1101/2020.06.21.163709

**Authors:** Yue Qin, Casper F. Winsnes, Edward L. Huttlin, Fan Zheng, Wei Ouyang, Jisoo Park, Adriana Pitea, Jason F. Kreisberg, Steven P. Gygi, J. Wade Harper, Jianzhu Ma, Emma Lundberg, Trey Ideker

## Abstract

The eukaryotic cell is a multi-scale structure with modular organization across at least four orders of magnitude^1,2^. Two central approaches for mapping this structure – protein fluorescent imaging and protein biophysical association – each generate extensive datasets but of distinct qualities and resolutions that are typically treated separately^3,4^. Here, we integrate immunofluorescent images in the Human Protein Atlas^5^ with ongoing affinity purification experiments from the BioPlex resource^6^ to create a unified hierarchical map of eukaryotic cell architecture. Integration involves configuring each approach to produce a general measure of protein distance, then calibrating the two measures using machine learning. The evolving map, called the Multi-Scale Integrated Cell (MuSIC 1.0), currently resolves 69 subcellular systems of which approximately half are undocumented. Based on these findings we perform 134 additional affinity purifications, validating close subunit associations for the majority of systems. The map elucidates roles for poorly characterized proteins, such as the appearance of FAM120C in chromatin; identifies new protein assemblies in ribosomal biogenesis, RNA splicing, nuclear speckles, and ion transport; and reveals crosstalk between cytoplasmic and mitochondrial ribosomal proteins. By integration across scales, MuSIC substantially increases the mapping resolution obtained from imaging while giving protein interactions a spatial dimension, paving the way to incorporate many molecular data types in proteome-wide maps of cells.

---

Advances in confocal microscopy and immunofluorescence (IF) imaging have created systematic pipelines for mapping the spatial distribution of proteins and other molecules within single cells^3,7,8^. Based on these techniques, the Human Protein Atlas (HPA) has launched an extensive effort to map protein subcellular locations using a library of specific fluorescent antibodies targeting more than 13,000 human proteins^5,9^. The use of multiple dyes with separate emission spectra enables locations to be determined relative to known landmarks such as the nucleus, cytoskeleton and endoplasmic reticulum, with the result that most human proteins can be assigned relative positions at sub-micron resolution.

In parallel, advances in mass spectrometry (MS) have provided a complementary means of mapping protein coordinates through their biophysical associations with other proteins^4,10^. MS is now routinely combined with affinity purification (AP-MS)^11^ or proximity-dependent labeling^12–16^ to enumerate physical protein-protein interactions *in vitro* or *in vivo*. Combining AP-MS with epitope tagging, the BioPlex project has generated a systematic map of physical interactions covering approximately 7,500 epitope-tagged human proteins (BioPlex 2.0)^6^, including approximately 56,000 candidate interactions organized into more than 400 multimeric protein complexes.

Given that protein imaging and biophysical association are leading approaches for mapping cell structure, with growing data resources and proteome-wide maps, a key question is whether and how they should be properly combined. One platform is sometimes used to validate results of the other, for example testing whether proteins associated by AP-MS co-localize within IF images^17^. The two platforms provide complementary measures of protein localizations, albeit at different physical scales, with IF positioning proteins globally relative to broad cellular landmarks, and AP-MS positioning proteins locally relative to other nearby proteins. In either case, such positioning has become increasingly quantitative in recent years, based on the ability of deep learning systems to recognize complex patterns in data^18–23^. Once an IF or AP-MS dataset has been analyzed to determine protein positions (or position distributions), a natural next step is to compute distances between these distributions, raising the possibility that the two types of protein distance might be calibrated and combined within integrated maps of cellular architecture (**Fig. 1a**).

**Fig. 1.**
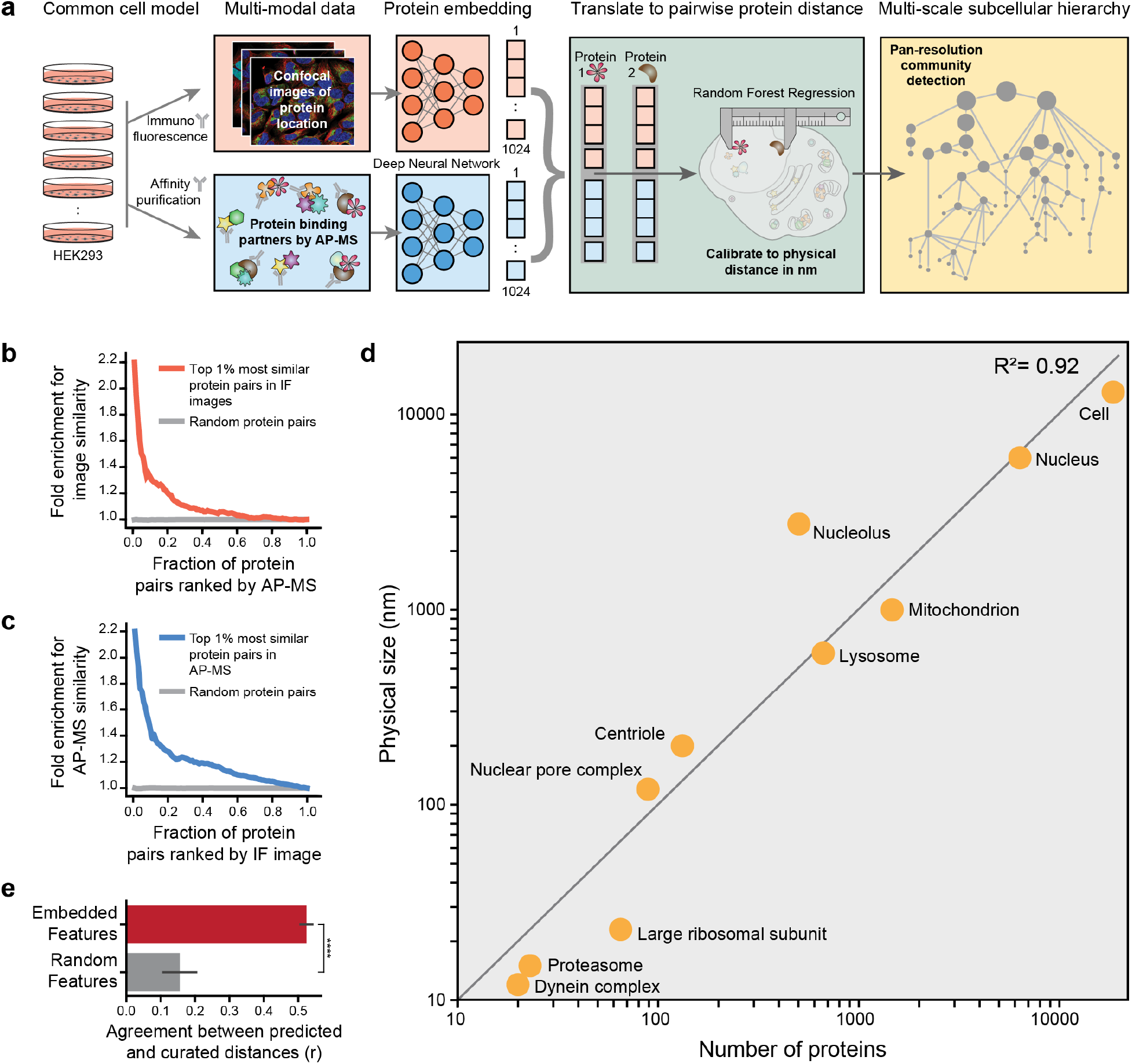
Fusing protein distances from immunofluorescence and affinity purification. **a,** Workflow. **b, c,** Protein pairs ranked by similarity in AP-MS data enrich for the most similar protein pairs in IF data (**b**), and vice versa (**c**). **d,** Calibrating size of subcellular components against the number of proteins assigned to the corresponding GO terms across four logs of physical scale. **e,** Performance of data-driven model in recovering protein-protein distances estimated using GO (red, Pearson *r*). Error bars show standard deviation in cross validation. Equivalent calculation for random feature sets (gray). ****, p < 0.0001.

## Protein position and distance, two ways

We assembled a matched dataset consisting of IF images from HPA^5^ and AP-MS data from BioPlex^6^. Both resources are partially based on HEK293 human embryonic kidney cells, leading us to 661 proteins with compatible imaging (1,451 images including replicates, **Extended Data Fig. 1**) and biophysical association data (291 proteins affinity-tagged as ‘baits’, the remaining 370 arising as interacting ‘preys’, **Supplementary Table 1**). While the number of studied proteins could be expanded by pooling additional IF and AP-MS datasets gathered in disparate cell types and conditions, we considered the importance of a controlled cellular context in prototyping any new approach. Using a deep convolutional neural network trained to recognize patterns in IF images^18^, we embedded each protein as a 1024-dimension feature vector, capturing its spatial distribution relative to counter-stained cellular landmarks (**Methods**). Similarly, the node2vec deep neural network^20^ was used to embed each protein as a second 1024-dimension feature vector based on its interaction neighborhood within the AP-MS data, including directly and indirectly associated proteins (**Methods**, **Extended Data Fig. 2**).

Next we computed protein-protein distances (cosine distance) for all pairs of proteins, separately in the IF and AP-MS high-dimensional embeddings. We found that the closest protein pairs measured by one technique were significantly enriched for those measured as close by the other, demonstrating that despite their differences, the two measurement types share significant information (**Fig. 1b, c**). As a means of calibrating the two distance measures, we sampled subcellular components of known physical size spread over several orders of magnitude in scale, from protein complexes of <20 nm to organelles >1 μm in diameter. The diameter of each component strongly correlated with its number of protein subunits documented in the Gene Ontology (GO)^24,25^, suggesting a general means of converting current GO annotations for a protein pair to an approximate distance in nanometers (**Fig. 1d**). Using these approximated distances as training examples, we taught a supervised machine learning model (random forest regression, **Methods**) to translate patterns in the concatenated IF and AP-MS features to an integrated measure of pairwise protein distance (**Fig. 1e**).

## A subcellular hierarchy at 20-20,000 nm

We analyzed this full set of protein distances to identify communities of proteins in close mutual proximity, suggesting distinct cellular components. Protein communities were identified at multiple resolutions, starting with those that form at the smallest protein-protein distances then progressively relaxing the distance threshold (multi-scale community detection^26,27^, **Methods**). Communities at smaller distances were contained, in full or in part, inside larger communities as the threshold was relaxed, yielding a structural hierarchy (**Fig. 2a**). The sensitivity of community detection was tuned for best concordance with two independent datasets not used elsewhere in our study: a separate collection of protein interactions reported in the Human Cell Map^28^ using proximity biotinylation, also in HEK293 cells, and patterns of gene co-essentiality observed in the Cancer Cell Dependency Map^29,30^ (**Methods**). This exercise also provided an end-to-end validation of our analysis pipeline (**Fig. 1a**), as the agreement with the outside datasets was significant over a wide range of community detection parameters (**Extended Data Fig. 3**). The final hierarchy, which we call the Multi-Scale Integrated Cell (MuSIC 1.0), contained 69 protein communities representing putative subcellular systems organized by 87 hierarchical containment relationships (**Fig. 2a**, **Supplementary Table 2**). Sixteen systems were contained in multiple larger ones, suggesting these systems have pleiotropic roles or are intermediate subassemblies of a larger complex. To elucidate the biological roles of each system, we aligned the MuSIC hierarchy to the equivalent literature-curated hierarchy provided by GO (**Methods**). A total of 46% of systems had significant overlap with GO; the remaining 54% did not clearly correspond to a known component and were labeled as putative novel (**Fig. 2a**). Where possible, systems were labeled by synthesizing prior literature with our own biological knowledge and reasoning.

**Fig. 2.**
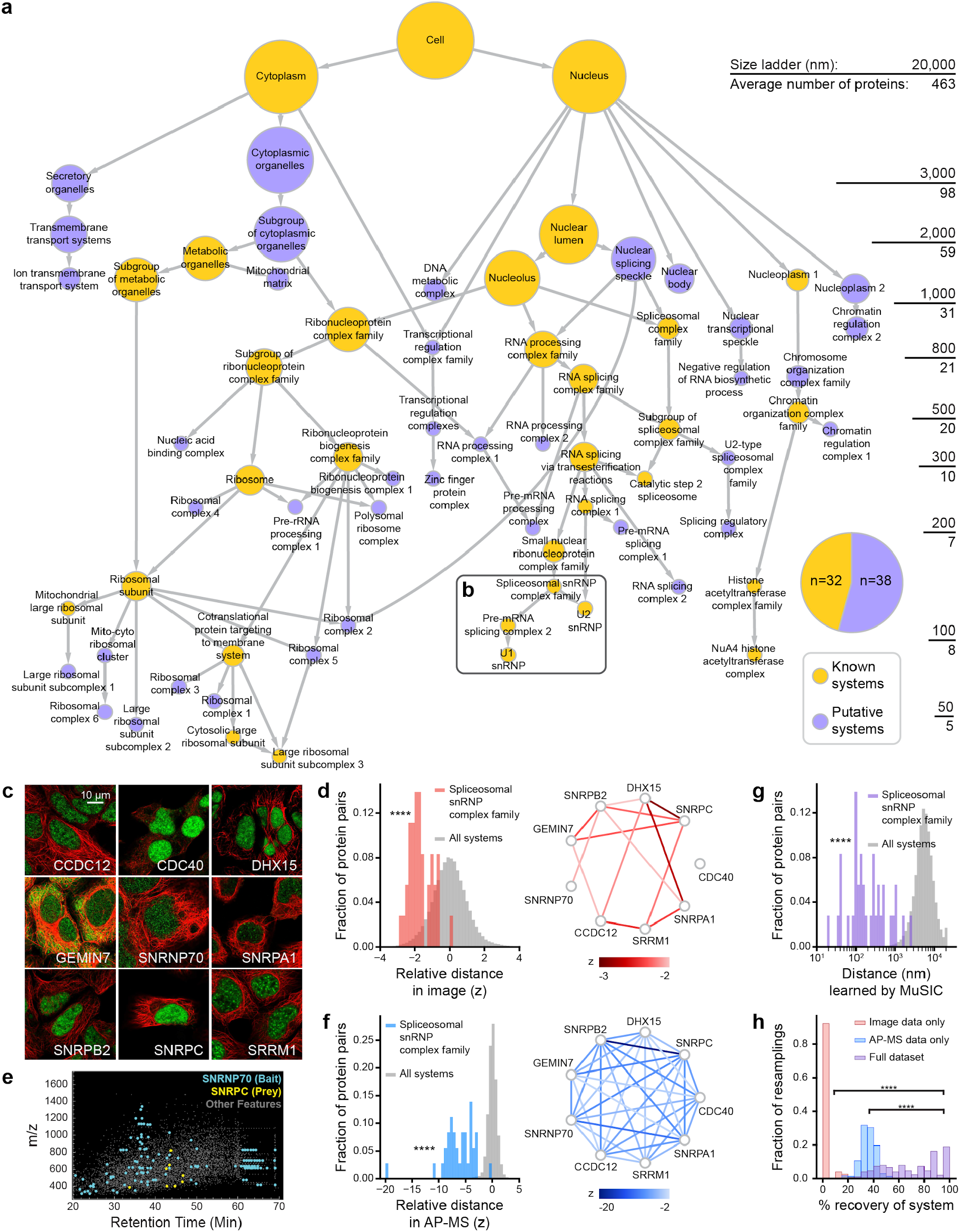
The Multi-scale Integrated Cell. **a,** MuSIC hierarchy, with nodes representing systems and arrows indicating containment of one system by another. Pie chart shows number of known (gold) or putative novel (purple) systems. **b,** Position of spliceosomal snRNP complex family. **c,** Immunostained spliceosomal snRNP complex proteins (green) or cytoskeleton (red). **d**, Corresponding IF protein-protein distances shown as *z*-scores (red), calibrated to all distances in IF data (gray). Results also shown as a network, with links for *z* ≤ −2, intensity showing magnitude of *z*. **e,** MS1 features identified during LC-MS analysis of a selected pull-down. Features corresponding to SNRNP70 (bait, cyan) and SNRPC (prey, yellow) are highlighted. **f,** Histogram and network as in (**d**), showing AP-MS rather than IF data. **g,** Integrated distances learned by MuSIC for spliceosomal snRNP complex proteins. **h,** Robustness analysis showing percent recovery of system when perturbing the input data; results accumulated over 300 resamplings (**Methods**). ****, p < 0.0001.

For example, MuSIC identified a community of nine proteins (labeled “Spliceosomal snRNP complex family”, **Fig. 2b**), the majority of which were specific members of the U1 and U2 small nuclear ribonucleoproteins (snRNPs) or had snRNP-related activities in previous studies^31–33^. Co-location of these proteins was supported by IF, which showed a common nucleoplasm signal (**Fig. 2c**) and corresponding close distances in the IF embedding (**Fig. 2d**). The image distributions varied considerably, particularly across nucleoplasm and cytoplasm, such that most but not all protein pairs were closely associated. Close proximities of these proteins were reinforced by the AP-MS data, however (**Fig. 2e, f**), based on similar proteome-wide patterns of interaction. The combined support from both data types induced this spliceosomal complex to form a specific protein community at a resolution of around 150 nm (**Fig. 2g, h**).

## Added sensitivity, resolution, context

We next examined the sensitivity with which these same systems could be identified using IF or AP-MS only, versus the full dataset. In general, we found that IF tends to robustly identify large systems such as organelles and sub-organellar parts that form the upper structure of MuSIC, whereas AP-MS robustly identifies smaller subcomponents within IF systems (**Fig. 3a, b**, **Methods**). For the majority of systems, the highest robustness was observed when integrating both types of data (**Fig. 3c, d**). For instance, the ability to recover the spliceosomal snRNP complex family dropped considerably when using only AP-MS data compared to integrated analysis (**Fig. 2h**).

**Fig. 3.**
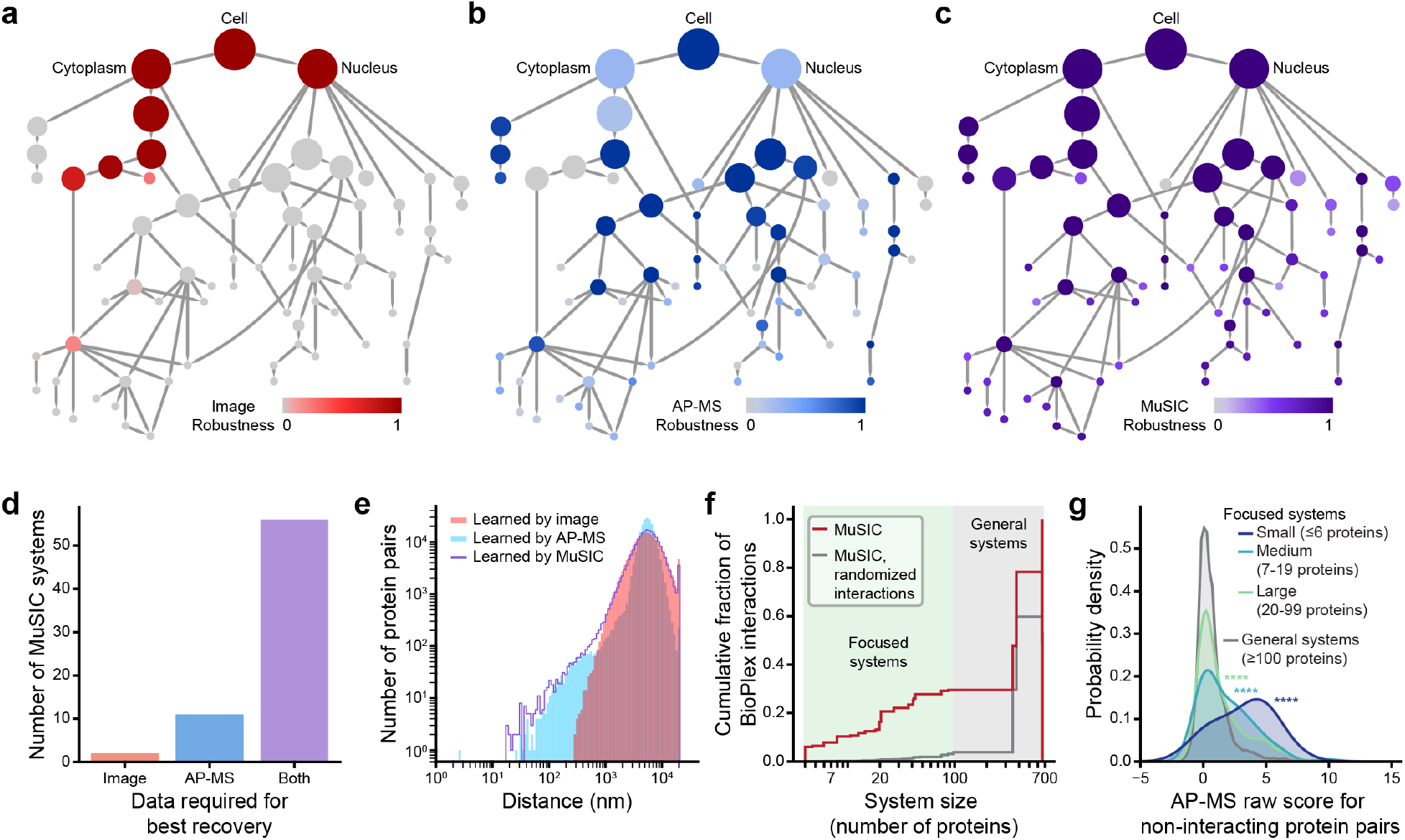
Different data, different scales of information. **a-c**, MuSIC hierarchy colored with system robustness using IF only (**a**), AP-MS only (**b**), or integrated data (full MuSIC, **c**). **d**, Number of systems for which the highest robustness was obtained with IF, AP-MS, or both data types (root system excluded). **e**, Distribution of protein-protein distances learned using only IF (red), only AP-MS (blue), or integrated analysis (purple). **f**, Cumulative fraction of BioPlex protein interactions within MuSIC systems (red) versus random protein pairs (gray, 1000 randomizations). **g**, Distribution of AP-MS raw z-scores for protein pairs not labeled as interacting by BioPlex. P-values calculated against general systems ≥100 proteins. ****: p < 0.0001.

Further insight came from examining the complete distributions of distances inferred using each data type separately (**Fig. 3e**). Using IF only, the measured protein distances were generally >200 nm with a median of approximately 5000 nm – the scale of large intracellular assemblies and organelles. Using AP-MS only, the median protein distance was also around 5000 nm, as expected if most proteins do not physically interact; however, a fraction of pairs were assigned extremely short distances of 20-200 nm – the realm of protein interactions and complexes. By integrating both data types, MuSIC distance measurements spanned four orders of magnitude in physical scale, ranging from roughly 20 to 20,000 nm (**Fig. 3e**).

Notably, 30% of AP-MS protein interactions fell within a focused system of <100 proteins (**Fig. 3f**). In each of these cases, such knowledge validates and aids in biological interpretation of the protein-protein association. We found that such context also allows rescue of protein interactions with support in the raw AP-MS spectra but which were overlooked in previous proteome-wide analysis due to the stringent scoring thresholds necessary to control for false discoveries. Among protein pairs not reported to interact in the previous BioPlex study^6^, pairs in smaller systems had significantly stronger AP-MS scores than pairs in larger systems (*p* < 0.0001, **Fig. 3g**), suggesting an untapped trove of bona-fide physical interactions.

## Exploration of newly identified systems

Of the 661 proteins common to the IF and AP-MS datasets, 370 had not yet been affinity tagged as the central baits of an AP-MS experiment – rather, they had appeared in the list of preys isolated by another affinity-tagged protein. Therefore, as an immediate means of validating candidate systems in MuSIC, we created affinity tags for 134 former preys and performed AP-MS, resulting in the identification of 339 physical interactions (**Supplementary Table 1**). We found that 44/69 MuSIC systems were specifically enriched for the new interactions (64%, FDR < 0.1, **Fig. 4a**), including 23 “putative” novel systems. Conversely, 195 (58%) of new interactions fell into MuSIC systems of <100 proteins, placing these into specific subcellular contexts (**Extended Data Fig. 4**).

**Fig. 4.**
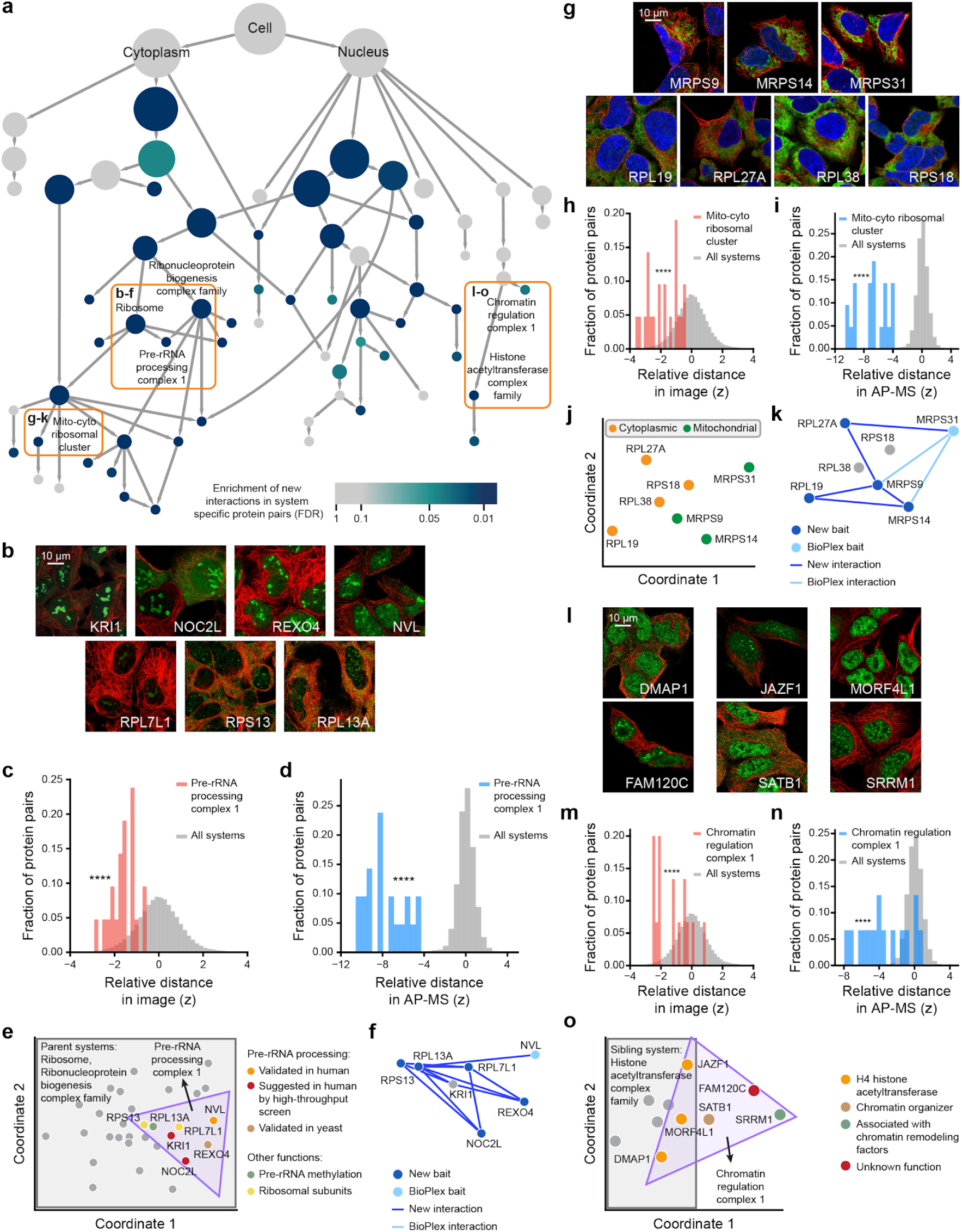
Validation of MuSIC systems using new AP-MS pull-downs. **a,** MuSIC hierarchy with system color showing enrichment for new AP-MS interactions at indicated false discovery rate (FDR, blue gradient, **Methods**). Orange boxes indicate systems highlighted in text. **b,** IF images showing similar nucleolar signals among proteins in the pre-rRNA processing complex 1. **c, d,** Corresponding distributions of protein-protein distance z-scores for IF (**c**, red) or AP-MS (**d**, blue), calibrated to all such distances, respectively (gray). **e,** Spring embedding of proteins in parent systems (grey box) of pre-rRNA processing complex 1 (purple triangle) based on pairwise distances learned by MuSIC. Proteins colored according to known functions. **f**, Validated physical interactions in pre-rRNA processing complex 1. **g-k**, Similar analyses for mito-cyto ribosomal cluster. **l-o,** Similar analyses for chromatin regulation complex 1. Image colors represent immunostained protein (green), cytoskeleton (red), or nucleus (blue). ****, p < 0.0001.

Among the novel systems validated by the additional interaction data was an assembly of seven proteins (**Fig. 4b-f**), which we named ‘Pre-ribosomal RNA (pre-rRNA) processing complex 1’ based on synthesizing information about known protein activities^34,35^ (NVL, RPL13A) with results from human high-throughput genetic screens^36^ (KRI1, NOC2L) and orthology to budding yeast^37^ (REXO4, **Fig. 4e**). The close association of these proteins was based on IF similarity (predominantly nucleolar localizations, **Fig. 4b, c**) and common AP-MS interaction neighborhoods (**Fig. 4d**), although, notably, none of the protein pairs had been reported to directly interact in BioPlex (**Fig. 4f**). Our new affinity purifications targeted five of these proteins as baits, recovering direct interactions for the majority of protein pairs in the system (11/21, **Fig. 4f**). We noted that the recovery of interactions was specific to that system (52.4% interactions recovered), in comparison to interactions with proteins in closely related systems (23.6%) or elsewhere in MuSIC (0.5%, **Extended Data Fig. 5**).

A second example was a system of seven proteins featuring abundant crosstalk between canonical subunits of the cytoplasmic and mitochondrial ribosomes (**Fig. 4g-k**). All showed cytoplasmic staining, explaining their close IF distances (**Fig. 4g-h**, staining is dim for MRPS9, MRPS14 and MRPS31 compared to their predominant mitochondrial locations). The IF similarity was further supported by AP-MS neighborhood similarity, although majority of the protein pairs had not yet been reported to physically interact (**Fig. 4i-k**). In the new affinity pull-downs, we tagged four of these proteins as baits (two cytoribosomal, two mitoribosomal), identifying five direct within-system physical interactions, four interconnecting cytoplasmic and mitochondrial factors (**Fig. 4k**). Such crosstalk has not been previously reported but may play a role in mitoribosome biogenesis, a poorly understood process^38^. Based on these findings, we named this system ‘Mito-cyto ribosomal cluster’.

A number of systems captured novel protein associations apart from those we had targeted with the new AP-MS data. One example (**Fig. 4l-o**) was a system of six proteins, of which DMAP1, MORF4L1, and JAZF1 were histone acetyltransferases^39,40^, SATB1 had been shown to interact with histone acetyltransferases^41^, SRRM1 had been implicated in chromatin remodeling^42^, and FAM120C had no previously assigned function (**Fig. 4o**). The specific association of these proteins was supported by their close measured distances in both IF (nucleoplasm and nuclear speckles, **Fig. 4l, m**) and AP-MS data (**Fig. 4n**). We thus named this system ‘Chromatin regulation complex 1’ based on the previously documented protein functions (**Fig. 4o**). The close associations of FAM120C, a candidate disease protein in autism spectrum disorder^43^, with proteins involved in regulation of chromatin suggests a novel function for this protein, consistent with the previous identification of other chromatin factors in autism^44^.

## Discussion

In classical image analysis of cells, the locations of proteins are identified in reference to a small panel of known markers for organelles. Here, MuSIC moves away from this closed library of organellar locations to an open platform in which both existing and new structures are identified *de novo* from structure in data. Numerous synergies between the IF and AP-MS data support the identification of 69 hierarchically-organized protein systems, many of which are robustly identified only when integrating both data modalities (**Fig. 3d**).

What about when the data disagree? While nearly a third of AP-MS interactions fall within the same focused system of <100 proteins, more than two thirds do not (**Fig. 3f**). For example, PPP6R1, a phosphatase, and NPAS1, a helix-loop-helix transcription factor, interact directly by AP-MS but were placed in different organelles in MuSIC related to their distinct image locations (PPP6R1, Cytoplasm; NPAS1, Nucleus; **Extended Data Fig. 6a**). In general, such discrepancies may indicate rare physical association of the proteins in a common compartment despite their more abundant locations in distinct others, as might be expected from pleiotropic roles or stages of protein maturation^45^. Alternatively, the divergent measurements may be explained by temporal dynamics, by which the two proteins interact transiently or periodically (e.g. cell cycle, circadian rhythms). Discrepancies may also derive from the fact that IF detects endogenous proteins whereas AP-MS detects over-expressed tagged proteins. Barring the above explanations, one can always suspect errors in data, e.g., cases in which the physical interaction represents spurious or non-specific binding. Some disagreement between IF and AP-MS can clearly be tolerated by the system, such as MuSIC’s correct assignment of GEMIN7 and SNRNP7 into the U1 small nuclear ribonucleoprotein particle^32,46^ (U1 snRNP, **Fig. 2a**), despite only partial overlap in the image distributions of the two proteins (**Extended Data Fig. 6b**). In this case, correct assignment was facilitated by the direct association of these proteins by AP-MS.

While the imaging field is accustomed to thinking about physical sizes and distances between cellular objects, the idea that protein interactions can be calibrated to measure physical distance is, to our knowledge, new to this study. The estimates here suggest that protein networks can measure sizes of cellular components down to about 20 nm (**Fig. 2a**). In contrast, the minimum size resolution learned for IF imaging, at approximately 200 nm (**Fig. 3e**), is in good agreement with conventional estimates^47^. This type of integrated analysis, with distance calibration, allows discovery of biological structure across a wide range of physical scales, including those scales that fall between the windows of resolution for each individual assay. It will be interesting to explore the size resolutions of other omics technologies, many of which might be calibrated to measure molecular distances and, in turn, contribute to integrated maps of the multi-scale cell.

## Acknowledgements

We gratefully acknowledge helpful discussion and comments from Cherie Ng, members of the Ideker laboratory, members of the Lundberg laboratory, the Human Protein Atlas, and the anonymous referees. This work was supported by the National Institutes of Health (NIH) under grants P41 GM103504 and R01 HG009979 to T.I., U24 HG006673 to E.L.H., S.P.G, and J.W.H., R50 CA243885 to J.F.K., and by the Knut and Alice Wallenberg Foundation (2016.0204) and the Swedish Research Council (2017-05327) to E.L..

## Author Contributions

Y.Q., E.L. and T.I. designed the study and developed the conceptual ideas. C.F.W. and W.O. generated image features. E.L.H., J.W.H., and S.P.G. generated AP-MS data. Y.Q. and J.M. designed the data integration approach. Y.Q. and F.Z. designed the community detection approach. All authors contributed in developing ideas for data analysis. Y.Q. implemented all computational methods and analyses. Y.Q. and T.I. wrote the manuscript with input from all other authors.

## METHODS

### Data sources

IF images interrogating protein locations in the HEK293 cell line were downloaded from the HPA Cell Atlas^5,18,19^. Physical protein interactions detected by AP-MS in the HEK293T cell line were downloaded from the BioPlex 2.0 protein interaction database^6^. We focused our study on the intersection of these IF and AP-MS datasets, i.e. selecting images of immunofluorescent proteins that had been affinity-tagged as baits or detected as preys in BioPlex. Each image had four channels, one for the protein of interest, and three for reference markers including nucleus, microtubules and endoplasmic reticulum. Images involving antibodies that targeted more than one protein, or which lacked annotated cellular localizations, were removed. The final imaging dataset contained 1,451 images covering 661 proteins and 726 antibodies, corresponding to a range of 2-6 images per protein (**Extended Data Fig. 1**). The final AP-MS dataset covered this same set of 661 proteins with 281 direct protein-protein interactions among the 661 proteins; we also retained the entire BioPlex 2.0 network of 10,961 proteins and 56,553 protein interactions, which provides significant information about the extended network neighborhoods of the 661 proteins (used in data embedding, see below).

### Data embeddings

Three different image embedding methods (DenseNet, SLF, Paired Cell Inpainting) were compared for their ability to enrich for the 281 BioPlex protein-protein interactions among the 661 proteins (**Extended Data Fig. 7a**). <u>DenseNet</u>: For each image, a 1024-dimension feature embedding was generated from a DenseNet-121 model^48^ optimized for annotating HPA images as previously described^18^. This method uses the protein channel and all three reference channels. SLF: Subcellular location features (SLF) were extracted at a per-cell level following the procedure described previously^19,49,50^. Here we used the same set of 719 features as in the previous SLF study^19^, capturing the relative relationship between the immunostained protein and the two major HPA cellular landmarks, nucleus and microtubules. <u>Paired Cell Inpainting</u>: The Paired Cell Inpainting^51^ unsupervised method uses a deep neural network to extract a 384-dimension feature vector for each protein, which was downloaded directly from http://hershey.csb.utoronto.ca/paired_cell_inpainting_features. For the SLF and Paired Cell Inpainting embeddings, per-image features were obtained by averaging the features of the cells within an image; the DenseNet embedding is already provided at the image level. We chose the DenseNet embedded features due to their ability to best enrich for physical protein interactions (**Extended Data Fig. 7a**).

For AP-MS data, three different embedding methods (node2vec, RWR, node properties) were compared for their ability to enrich for the 281 protein pairs with highest cosine similarity calculated in the IF images using the DenseNet embedding (**Extended Data Fig. 7b**; the number 281 was selected to match the number of protein-protein interactions used above for the reciprocal analysis). All three embedding methods input a protein-protein interaction network, in which nodes represent proteins and edges represent measured protein-protein interactions drawn from all available BioPlex AP-MS data (*n* = 10,961 nodes, *m* = 56,553 edges, see above). <u>Node2vec</u>: This method uses a deep neural network to learn feature representations of the network neighborhood surrounding each node^20^. A 1024-dimension feature embedding for each protein was obtained by applying node2vec with parameters *p* = 2, *q* = 1. <u>RWR</u>: Random walk with restart (RWR) is carried out according to equation:

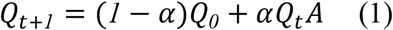

α tunes the distance that a protein signal diffuses within the protein-protein interaction network, here set to α = 0.6, which was determined to be the optimal setting in a previous human protein network study^52^. **Q**_*0*_ is an *n* ⨉ *n* identity matrix, and *A* is the column-normalized adjacency matrix representation of the network^53^. The embedding of each node was taken as the corresponding row of *Q_t_* if eqn. (1) converged or after 30 iterations. <u>Node properties</u>: Each protein embedding was represented by a short feature vector formulated from the following graph-theoretic properties: (i) degree, (ii) clustering coefficient, (iii) closeness centrality, (iv) normalized betweenness centrality, (v) eigenvector centrality, (vi) stress centrality, (vii) bridging centrality, and (viii) information centrality, following definitions from a previous study^54^. Node2vec embeddings were chosen based on superior performance in enriching for protein pairs with similar image embeddings (**Extended Data Fig. 7b**).

### Data integration

Protein annotations from the Cellular Component branch of the Gene Ontology (GO)^25^ were used to formulate a training set of approximate protein pairwise distances. GO was downloaded on 25.9.2018 from http://geneontology.org. Annotations based on HPA images were removed from GO to avoid circularity. Among the 661 proteins under study, 602 had specific GO annotations (i.e. other than the cellular_component root term) which were used to supervise the calibration of distance from embedded features. The relative GO similarity (converse of distance) for a protein pair *p_1_* and *p_2_* was calculated according to a previously proposed metric^55^ as:

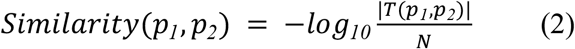

where |*T*(*p_1_*, *p_2_*)| is the number of proteins in the smallest term to which *p_1_* and *p_2_* are both annotated and *N* is the total number of proteins with specific annotations in GO.

Random forest regression^56^ implemented in the Python scikit-learn package^57^ was used to learn the approximate GO similarity, scaled to [0, 1], between a pair of proteins. The training features included the complete image and AP-MS embeddings of each protein in the pair as well as a panel of similarity measurements (cosine, Pearson, Spearman, Kendall, Manhattan, and Euclidean) of the protein pair in the image and AP-MS embeddings respectively. Because every protein had multiple images (ranging from two to six, see **Data sources** above), six different groups of image-to-protein mappings were randomly generated while requiring each image for a protein to be mapped at least once. We trained random forest regression with five-fold cross validation for each group. The final similarity of each protein pair (henceforth called the *<u>integrated protein-protein similarity network</u>*) was obtained by averaging the random forest predictions from the six groups. As a negative control, a 1024-dimension random vector sampled from a normal distribution was generated for each image embedding and each AP-MS embedding (**Fig. 1e**).

### Pan-resolution community detection

The integrated protein-protein similarity network was analyzed to detect distinct communities of similar proteins using the Clique eXtracted Ontology algorithm^27^ (CliXO v1.0, https://github.com/fanzheng10/CliXO-1.0). CliXO finds the maximal cliques in a weighted network while progressively decreasing the threshold on edge weights. Lower thresholds yield cliques that may contain, in full or in part, cliques identified at higher thresholds, resulting in a hierarchy of communities interrelated by community containment relations. CliXO has four parameters that control the depth (*α*), width (*β*), modularity (*m*) and modularity significance (*z*) of the community hierarchy (**Extended Data Fig. 3a**). We swept through 500 different combinations of these four parameters to obtain a pool of hierarchies. All communities, representing putative cellular systems, were required to have at least four proteins to further ensure quality and validity.

To select an optimal hierarchy, each hierarchy was evaluated based on its concordance with two independent datasets, the Human Cell Map^28^ and the Cancer Cell Dependency Map (v18Q2)^29^, which were not used elsewhere in this study (**Extended Data Fig. 3b, c**). From the Human Cell Map, 178 protein-protein interactions detected in HEK293 cells with high-confidence (FDR ≤ 0.01) were obtained, covering 293 proteins in MuSIC. From the Cancer Cell Dependency Map (DepMap)^29^, we selected 9,030 protein pairs, covering 629 proteins in MuSIC, for which the CRISPR gene disruptions of the two proteins led to significantly correlated fitness profiles across the panel of 436 DepMap cell lines (so-called “co-essential” proteins). For each hierarchy, we recorded the number of Human Cell Map protein-protein interactions (*x*) or DepMap co-essential protein pairs (*y*) that were covered by systems that were significantly enriched (FDR ≤ 0.1) for those interactions (**Extended Data Fig. 3b, c**). The hierarchy having the highest number (*x* ⨉ *y*) was selected for further study; among the several ties, we selected the hierarchy with the least number of systems (i.e. guided by the principle of parsimony).

The hierarchical structure was further matured by assigning additional hierarchical parent-child containment relations between pairs of systems having a containment index ≥0.75 and removing redundant systems having Jaccard index ≥0.9 with parent systems. The containment and Jaccard indexes between two systems *S_1_* and *S_2_* were calculated based on the following formulae:

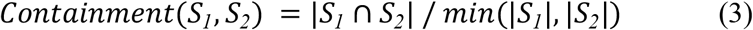

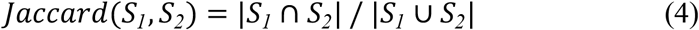

with |*S*| representing the number of proteins in a system. Redundant systems having only one protein difference from a parent system were also removed. When removing a system, the integrity of the hierarchical structure was maintained by adding containment relations from all children of that system to all of its parents.

We also analyzed the integrated protein-protein similarity network using Louvain clustering^26,58^ (https://github.com/vtraag/louvain-igraph, v0.6.1) which partitions network nodes into a set of distinct clusters. We ran Louvain 1,000 times and selected the partition that maximized the global modularity, as output by the algorithm. The partition used for the main MuSIC model included just two clusters, which were strongly enriched for proteins with known subcellular locations in the cytoplasm versus nucleus, respectively (**Fig. 2a**). These clusters were added as parent systems of the top-layer systems found by CliXO (described above). In particular, a CliXO system was added as a child of a Louvain system if at least 50% of its proteins were in the Louvain system(s). A similar process was employed to construct the jackknifed hierarchies in the robustness analysis (**Fig. 3a-c**, see **Dependence of systems on data types** below). For systems highlighted in main figures (**Figs. 2a, 4a**), we introduced a further quality control step in which we manually inspected the corresponding IF images and raw AP-MS spectra. This process prompted us to remove the protein RPL6 from the system “Mito-cyto ribosomal cluster” out of concerns for antibody correctness.

To label MuSIC systems as “known” or “putative” (**Fig. 2a**), MuSIC was aligned to the Cellular Component branch of GO, filtered for the proteins under study. A system *S* was considered “known” if there existed a GO term *T* that was significantly enriched (FDR ≤ 0.001, hypergeometric statistic) for proteins in *S* with *Jaccard* (*S*, *T*) ≥ 0.4, or if *Jaccard*(*S*, *T*) = 1, representing perfect agreement regardless of significance.

### Calibration to physical distance

The following calibration function was obtained using linear regression on the list of subcellular compartments with known physical sizes (**Fig. 1d**):

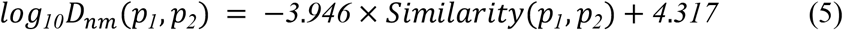

where *D_nm_*(*p_1_*, *p_2_*) is the physical distance in nm between proteins *p_1_* and *p_2_*, and *Similarity* (*p_1_*, *p_2_*) is calculated using eqn. (2) based on the number of proteins annotated to the compartment. The physical size of each MuSIC system (**Fig. 2a**) was calibrated based on the similarity threshold at which the system was found.

### Dependence of systems on data types

To investigate the relative contributions of IF and AP-MS data in system formation, we evaluated the robustness of each system under perturbations to the integrated protein-protein similarity network (**Figs. 2h, 3a-c**). To assess the dependence of each system on imaging data, we created an alternative network using IF features only, with AP-MS features randomized (see **Data integration** above). Subsequently, 10% of the edges in this protein network were randomly removed, and community detection was performed to construct a hierarchy using the same parameters as MuSIC (see **Pan-resolution community detection** above). This randomization procedure was repeated 300 times to obtain a pool of perturbed hierarchies, similar to statistical jackknifing^59^. The “percentage recovery” of a MuSIC system *S_MuSIC_* in a perturbed hierarchy *H_P_* was calculated as:

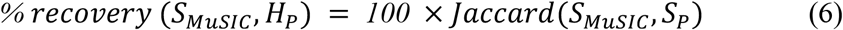

where *S_P_* is the perturbed system best enriching for *S_MuSIC_* in *H_P_* (**Fig. 2h**). *S_MuSIC_* was considered to be “recovered” by *H_P_* if the enrichment of *S_MuSIC_* in *S_P_* was significant (FDR ≤ 0.001, hypergeometric statistic) and *% recovery*(*S_MuSIC_*, *H_P_*) ≥ 40%, or if *% recovery*(*S_MuSIC_*, *H_P_*) = 100%, representing perfect agreement regardless of significance. The image robustness score for *S_MuSIC_* (**Fig. 3a**) was the fraction of all perturbed hierarchies that recovered *S_MuSIC_*.

To assess the dependence of each system on AP-MS data, a reciprocal procedure was performed by generating an alternative integrated protein-protein similarity network using AP-MS features only, with imaging features randomized (**Fig. 3b**). As a benchmark, we also computed the robustness of the full MuSIC model using the original integrated protein-protein similarity network without IF or AP-MS feature randomization (**Fig. 3c**), using the same jackknifing procedure described above.

### Global system validation using new AP-MS data

We constructed stable HEK293T cell lines for 134 bait proteins (**Supplementary Table 1**) with C-terminal FLAG-HA-tags based on the human ORFeome v8.1 (http://horfdb.dfci.harvard.edu/)^60^ as previously described^6,61,62^. Cell pellets were lysed using 50 mM Tris-HCl pH 7.5, 300 mM NaCl, 0.5% (v/v) NP40 buffer, and cell debris were removed with centrifugation and filtration. Mouse monoclonal anti-HA agarose resins (Sigma-Aldrich, clone HA-7), immobilized and pre-washed, were incubated with cell lysates at 4 ℃ for 4 hours. After removing supernatant, precipitates were washed four times with lysis buffer and two times with PBS (pH 7.2). Elution was performed in two steps by adding 250 μg/mL HA peptide in PBS at 37°C followed by TCA precipitation. Eluted samples were analyzed by LC-MS using Q-Exactive mass spectrometers (Thermo Fisher). Each bait protein was analyzed with biological duplicates for the affinity purification step and technical duplicates for the LC-MS step, yielding four replicates in total. MS/MS spectra were analyzed using the Sequest algorithm^63^ to match peptide sequences from the Uniprot database^40^ supplemented by signatures for green fluorescent protein (negative control), the FLAG-HA-tag, and common contaminants. Identified peptides and proteins were further filtered using the target-decoy method^64^ to control FDR. High confidence protein interactions were identified using the ComPASS algorithm^65,66^ on merged technical duplicates, followed by ComPASS-Plus analysis^62^.

To examine per-system enrichment for the new AP-MS data in MuSIC, the assignment of proteins to systems was shuffled while keeping the overall hierarchy structure and number of proteins per system the same, resulting in 1,000 random hierarchies. For each system, we calculated the empirical p-value^67^ for the number of new AP-MS interactions among the “system-specific” protein pairs, defined as protein pairs in that system but not in children of that system. P-values were corrected using Benjamini-Hochberg multiple test correction to obtain an FDR for per-system enrichment (**Fig. 4a**).

### Statistical analyses

All statistical tests were performed using SciPy^68^ with Benjamini-Hochberg multiple test correction as appropriate. Statistics involving comparison between two groups of data to assess differences in data distributions were calculated using either a Mann-Whitney U test, when samples were not paired (**Figs. 2–4**), or a Wilcoxon signed-rank test for paired samples (**Fig. 1e**). Statistics for assessing the enrichment of proteins or protein pairs were calculated using hypergeometric tests (**Extended Data Fig. 3**) unless stated otherwise.

### Data Availability

A web portal is available at http://nrnb.org/music with links to all major resources used for this study. These include the MuSIC systems map; the IF (HPA) and AP-MS data (BioPlex 2.0) on which the map is based; and data for the AP-MS pulldown experiments performed as follow-up. The follow-up AP-MS data have also been included as part of the larger compendium of protein interactions to be included in the next version of the BioPlex resource (BioPlex 3.0, *manuscript in review*).

### Code Availability

Links to code for performing feature embedding and multi-scale community detection are available at http://nrnb.org/music.

## EXTENDED DATA

**Extended Data Fig. 1.**
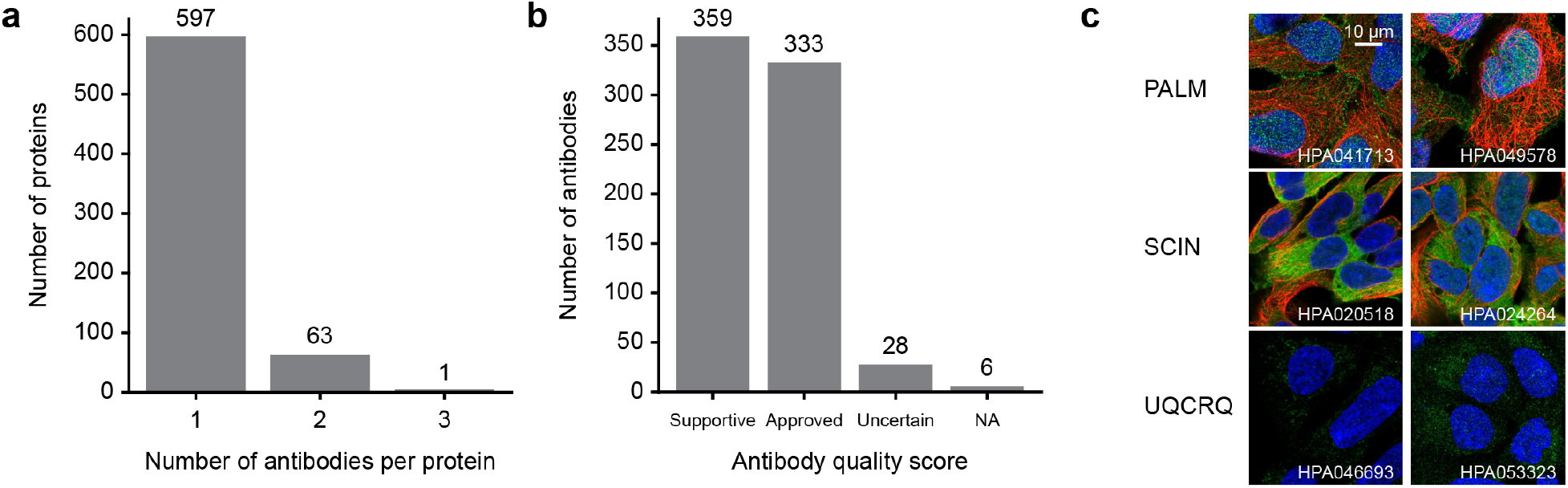
Antibody count and quality. **a,** Histogram showing the distribution in number of antibodies per protein over the proteins included in this study. **b,** Histogram showing the distribution in antibody quality scores over the antibodies used in this study. **c,** IF images for three pairs of antibodies targeting the same protein, respectively. Colors represent immunostained protein (green), cytoskeleton (red), or nucleus (blue). The images show high reproducibility even when different antibodies for the same target protein are used.

**Extended Data Fig. 2.**
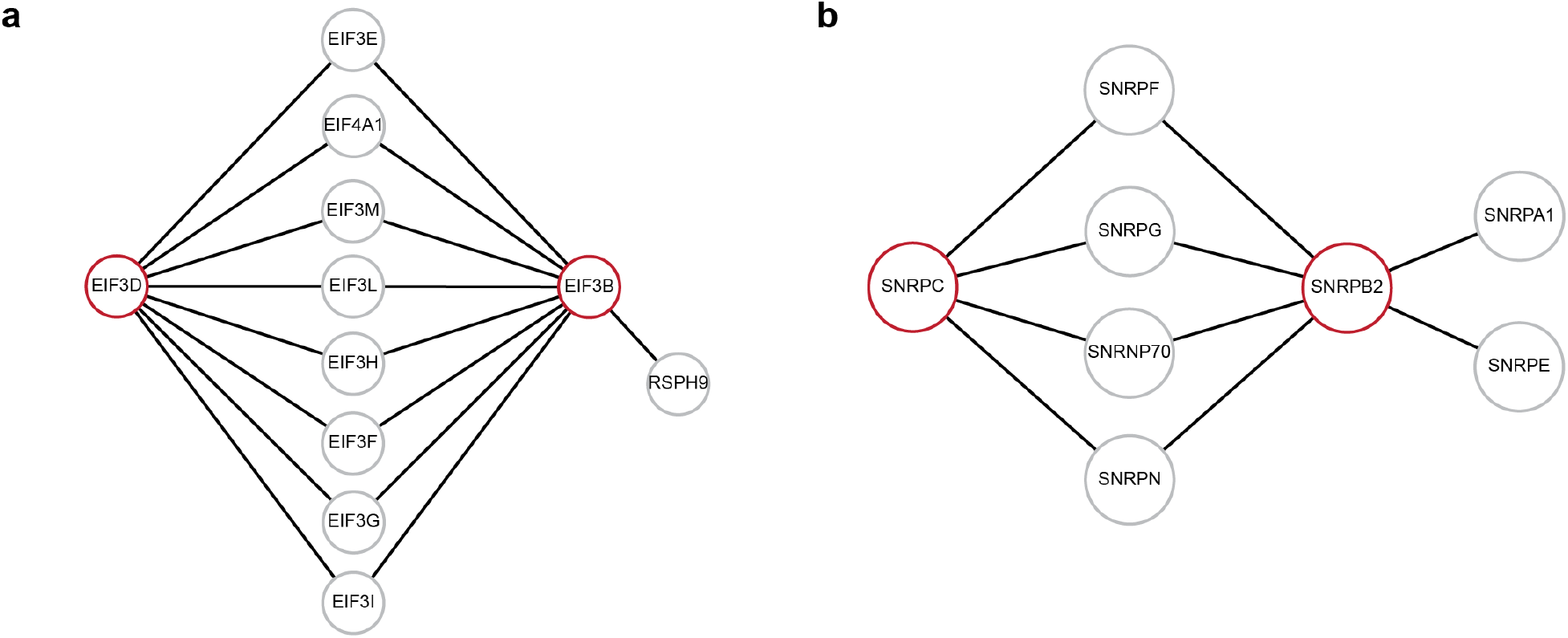
Non-interacting protein pairs nonetheless assigned high similarity in the AP-MS embedding. The node2vec network embedding^20^ can determine that two proteins have high similarity even if these proteins have not yet been recorded to have direct physical interaction. **a,** Network showing all proteins that physically interact with EIF3D and EIF3B (red) in BioPlex 2.0. EIF3D and EIF3B do not physically interact, but the cosine similarity of their embedded node2vec features, which consider the first interaction neighborhood (shown) and beyond (not shown), is 0.95. The maximum cosine similarity is 1. **b,** Network showing all proteins that physically interact with SNRPC and SNRPB2 (red) in BioPlex 2.0. SNRPC and SNRPB2 do not physically interact, but the cosine similarity of their embedded features is 0.93. In many cases where two proteins have high node2vec similarity, but do not directly interact, we found that neither protein had yet been tagged as bait for an affinity purification experiment. As the direct interaction has not yet been interrogated, the node2vec embedding serves to fill gaps in existing data. None of the proteins highlighted in red were tagged as bait proteins in BioPlex 2.0.

**Extended Data Fig. 3.**
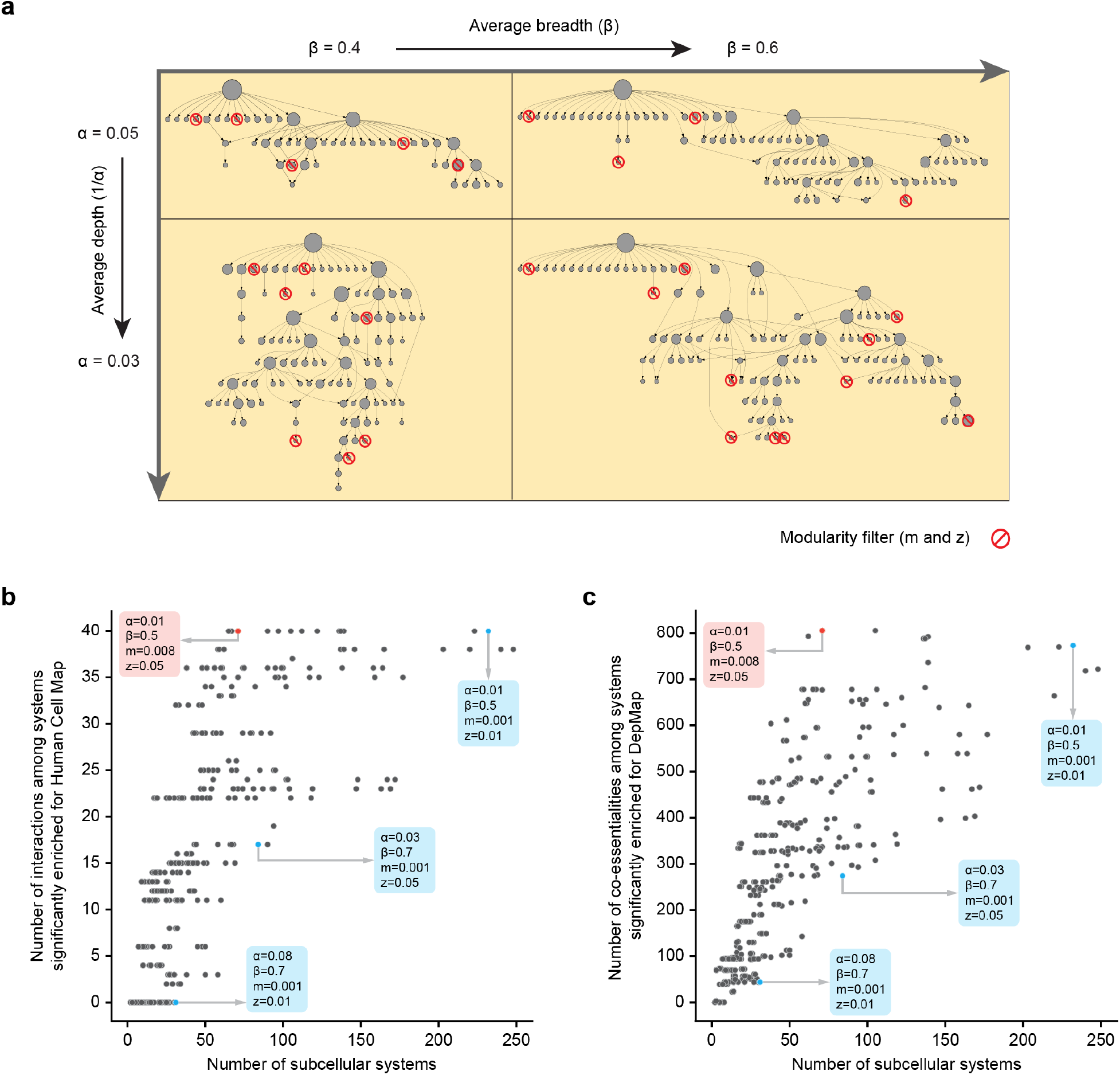
Selection of parameters for community detection. **a,** CliXO community detection has several parameters (breadth, x-axis; depth, y-axis; minimum modularity, red circle backslash) that affect the sensitivity with which communities are identified and thus the size of the hierarchy. **b, c,** Dotplots in which each dot is a hierarchy generated with a particular set of parameters. The selection for MuSIC is highlighted in red. This selection was among several that were optimal, based on enrichment for protein-protein interactions in Human Cell Map (**b**) and co-essentialities from DepMap (**c**). Examples of other parameters are shown in blue.

**Extended Data Fig. 4.**
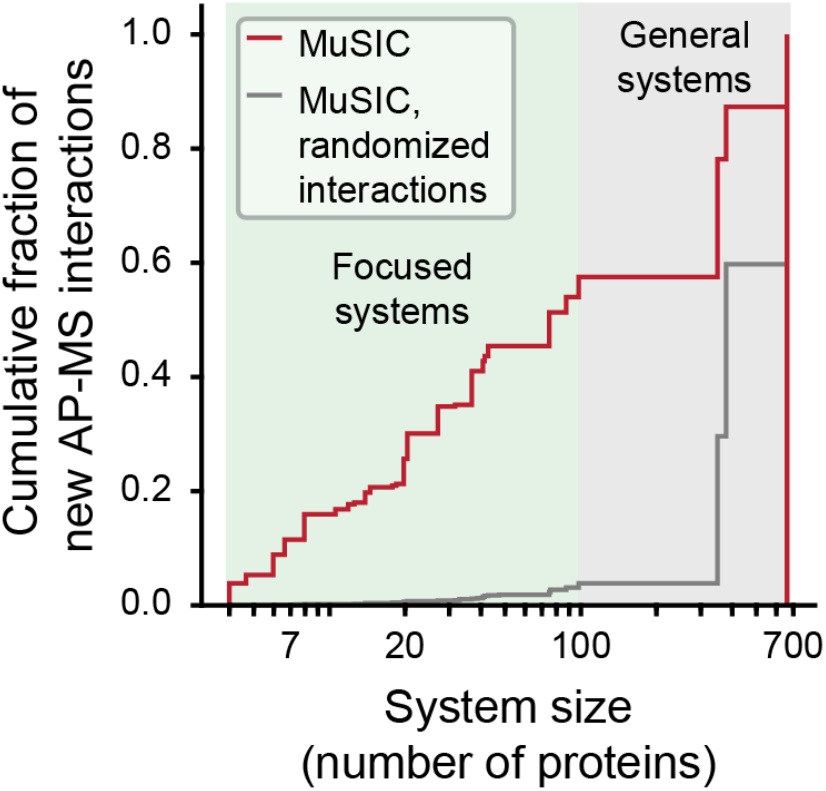
Cumulative fraction of new AP-MS interactions in MuSIC. Related to Fig. 3f.

**Extended Data Fig. 5.**
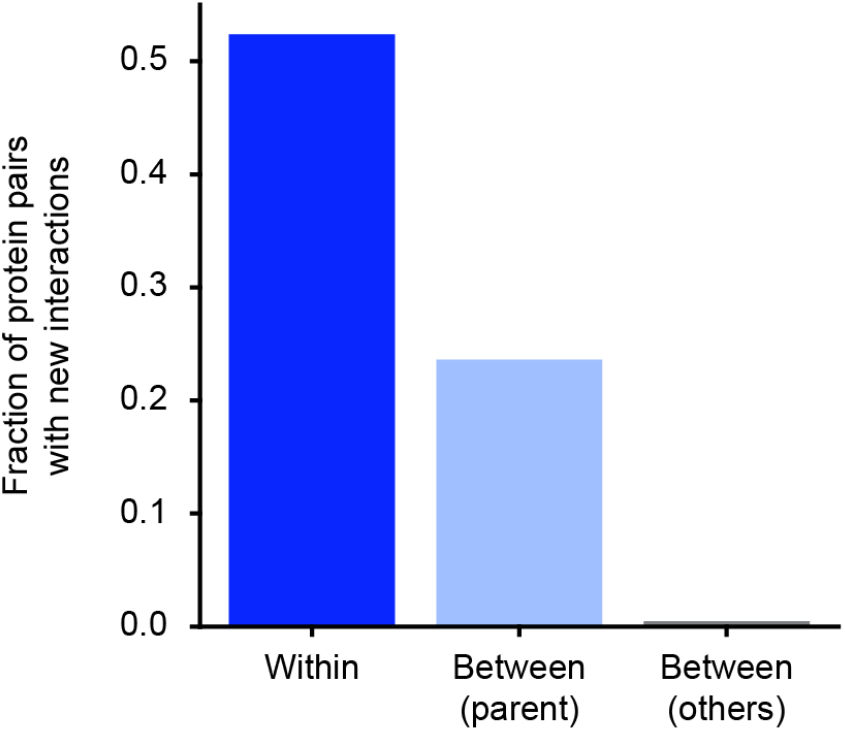
Classification of newly identified physical interactions for proteins in the pre-rRNA processing complex 1. The specific recovery of interactions within that same system is shown (dark blue bar), in comparison to interactions with other proteins organized under the same parent systems in the MuSIC hierarchy (ribosome, ribonucleoprotein biogenesis complex family, light blue bar) or with proteins organized elsewhere in MuSIC (gray bar).

**Extended Data Fig. 6.**
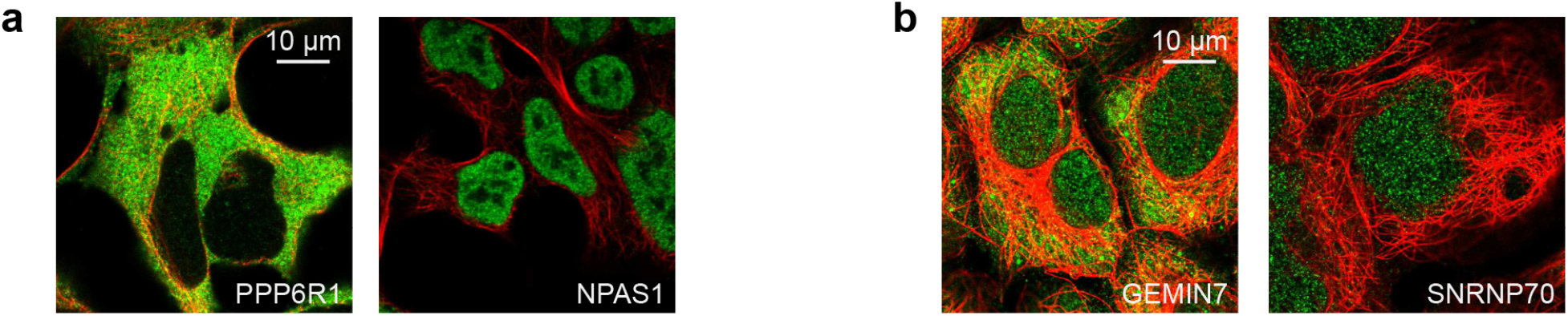
Examples of proteins with strong AP-MS protein interactions that have very different IF localization patterns. **a,** PPP6R1 and NPAS1. **b,** GEMIN7 and SNRNP70. Colors represent immunostained protein (green) and cytoskeleton (red).

**Extended Data Fig. 7.**
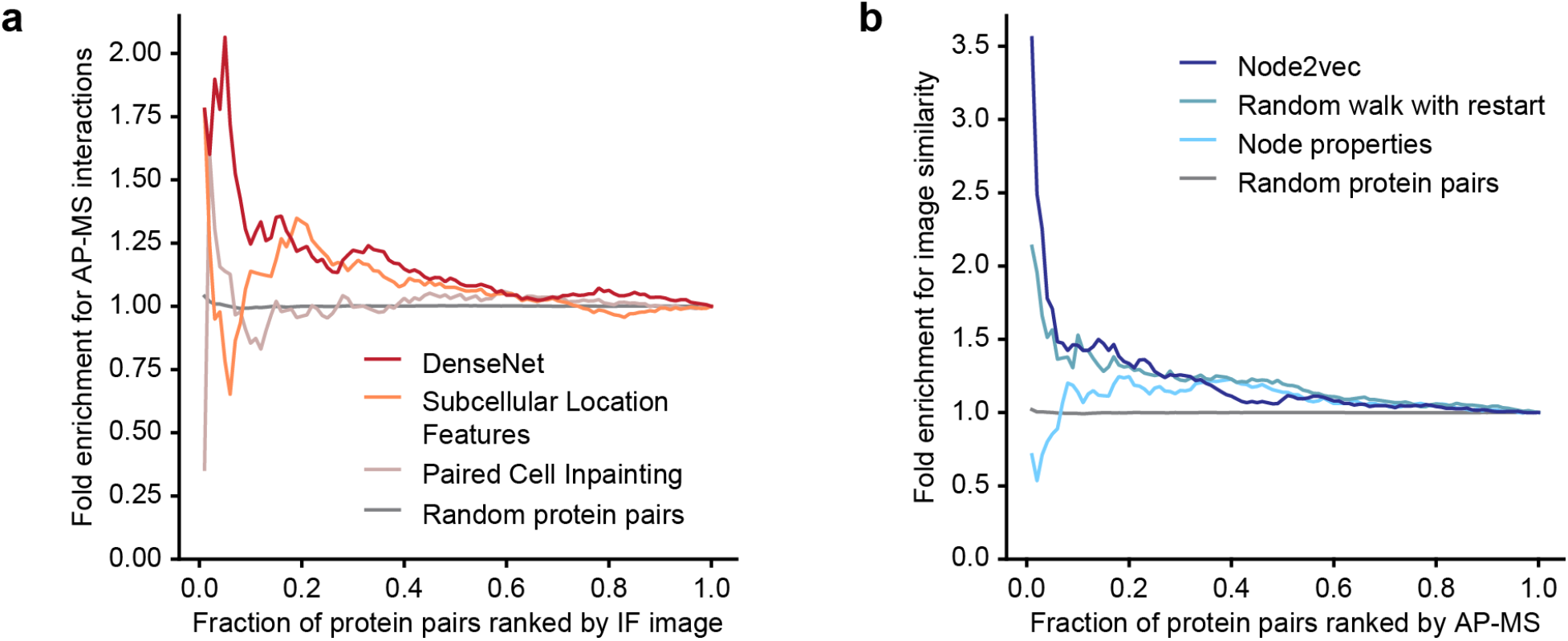
Comparison of different data embedding methods. **a,** Ability of different image embedding methods (colored curves) to generate image-image similarities (cosine similarity) in agreement with the set of all 281 protein-protein interactions in BioPlex. **b,** Ability of different AP-MS embedding methods to generate protein-protein similarities (cosine similarity) in agreement with protein pairwise similarities computed from HPA images (considering the top 281 protein pairs by image cosine similarity). Embedding methods are described in **Methods**.

